# Statistical analysis of number of genes and chromosome lengths of different microbial species

**DOI:** 10.1101/2022.08.13.503871

**Authors:** Wenfa Ng

## Abstract

Genome architecture concerns the organisation of genes on a chromosome, and has important implications to the fidelity in which genes are encoded on the chromosome, and how the information is read by DNA polymerase and RNA polymerase. This facet of genomics did receive attention in the early epoch of genomics, but it has received less attention in contemporary genomics as attention shifts to structural and functional genomics with the goal of annotating the function of each gene in the genome. This work sought to uncover relationships between number of genes and chromosome length in a variety of bacteria and archaea species as a preamble to understanding the prevalence and importance of repetitive sequences in the genome of prokaryotic species. Aggregate results with the ensemble of prokaryotic species profiled revealed a positive linear correlation between number of genes and chromosome length. Upon dissection into the Bacteria and Archaea domains, the linear relationship described above still stands for Bacteria but starts to break down in Archaea. This suggests that repetitive sequences are more important to Archaea species, which generally have a smaller genome (1.8 to 2.8 Mbp) and fewer genes (1500 to 2400) compared to bacterial species. In comparison, the bacterial genome is larger (4 to 5.6 Mbp), and encodes more genes (3600 to 5100). Overall, the results highlight that bacterial genome are efficiently encoded with few repetitive sequences. This, however, is not true for archaeal genome, which provides another line of evidence supporting the notion that archaea are ancestral eukaryotic cells, which like the archaea also houses large repetitive sequences.

**Graphical abstract:** 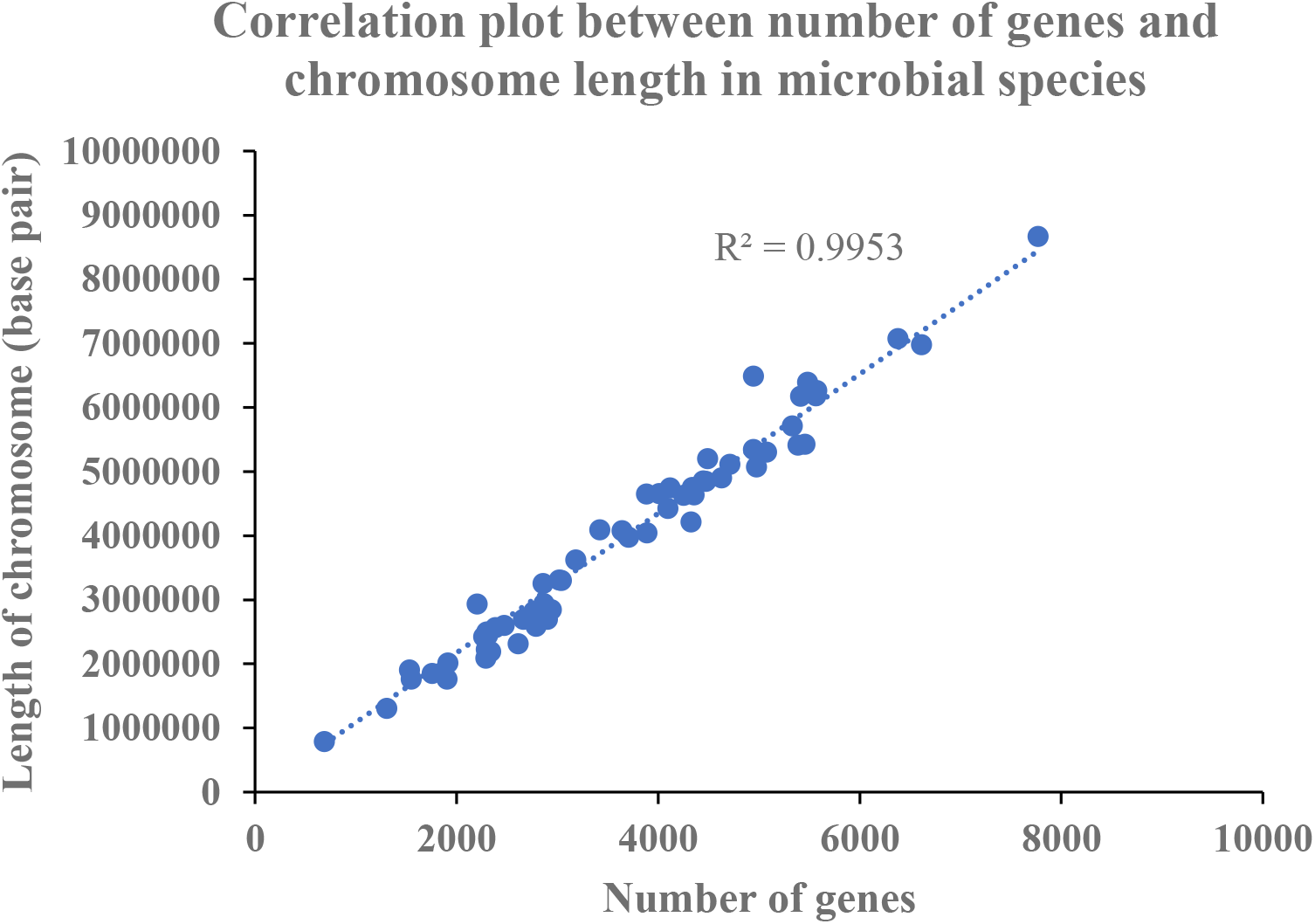

**Short description:** Statistical analysis across an ensemble of 59 microbial species revealed a strong linear correlation between number of genes and chromosome length. This suggests that prokaryotic genomes are highly compact with genes, and do not carry significant amounts of repeats unlike the case in eukaryotic organisms. The result holds significant implications for our understanding of genome evolution and compaction in prokaryotic organisms, and what drove their accession as foundational species of many ecosystems.

**Subject areas:** genomics, molecular biology, evolutionary biology, bioinformatics, systems biology,

## Introduction

Contemporary genomic studies typically focus on understanding the roles of genes in manifesting a phenotype as well as how regulatory processes are encoded in the genome in DNA sequence motifs. Such studies in functional genomics have yield a rich vein of information and knowledge that guides us in understanding and treating diseases as well as understanding physiological processes in the microbial world and ecosystem. More recently, the focus of genomics has partially turned to structural genomics in understanding how the shape and conformation of the DNA molecule encoding the genes help potentiate or regulate gene expression [1] [2]. While emerging in importance, structural genomics nevertheless touch on the question of how the genome is organised and regulated at the molecular level [3] [4].

This work attempts to answer a basic question concerning the relationship between the number of genes and genome length in the microbial world. Focussing on prokaryotic species in the domain Bacteria and Archaea, the study uses an in-house MATLAB encoded genome analysis software to retrieve information on the number of genes and genome length from each microbial species. This information is subsequently aggregated and used to plot a scatter plot that hopefully reveal the correlation between number of genes and genome length in the prokaryotic world as well as the domain level. To help further distill the range of values for number of genes and genome length at the level of Bacteria and Archaea, histogram plots were used to help identify the most prevalent range of values for the above two parameters of significance to the genomics researcher.

## Materials and Methods

Genome sequence of different species of bacteria and archaea were downloaded from Genbank, and processed with an in-house MATLAB genome analysis software to extract information on number of genes and genome length. Data from different species were aggregated and plotted as a scatter plot between number of genes and genome length. To discern if a linear relationship could be obtained between number of genes and genome length for different domains of life, individual scatter plots featuring the above two variables were completed for species in Bacteria and Archaea. In addition, histograms attempting to discern distribution of genome length and number of genes in different bacteria and archaea species were also plotted.

## Results and Discussion

Scatter plots are widely used in engineering and the natural sciences to reveal correlation or relationships between two variables postulated to regulate each other. Hence, x-y scatter plot was used to reveal whether number of genes and genome length are positively correlated in the Bacteria and Archaea domains. Figure 1 shows the result of this correlative plot, and the data shows that in the Bacteria domain, there is a strong positive linear correlation between number of genes and genome length across many bacterial species with correlation coefficient of 0.9967. In addition, there is likely a constant relationship between length of repetitive sequence and gene length in many bacterial species. More importantly, the data suggests that repetitive sequences that bloated the size of the genome in many species is less likely to be a problem in bacterial species. Hence, bacterial genome can be characterised as efficiently encoded with little sequence space taken up by repetitive sequences such as is the case for many eukaryotic genomes.

**Figure 1:**
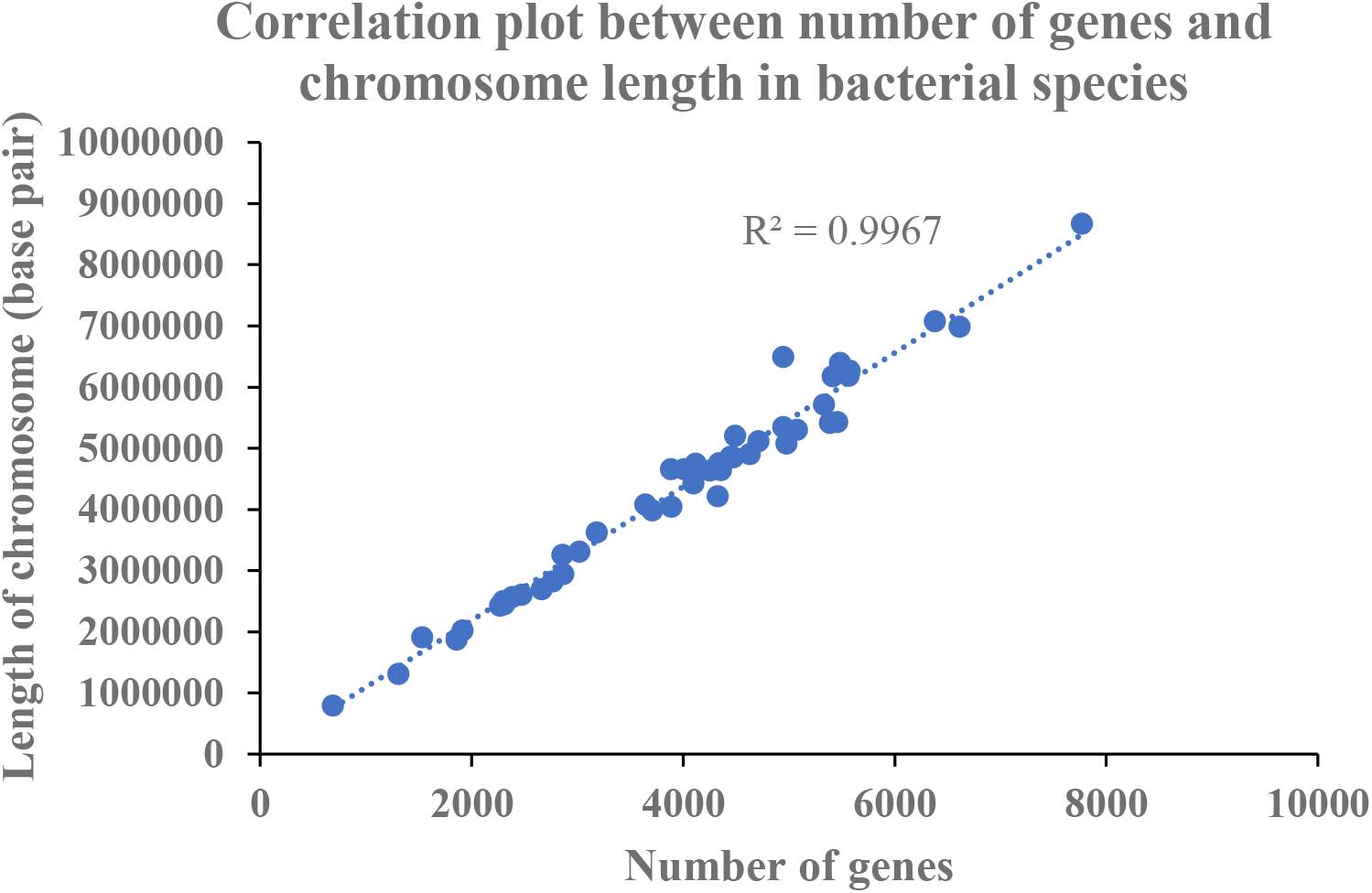
Correlation plot between number of genes and chromosome length in bacterial species. The data shows that there is a good linear correlation between number of genes and length of chromosome across the Bacteria domain, indicating that repetitive sequences are not highly pervasive in the bacterial species

Figure 2 and 3 shows the distribution of number of genes and chromosome length in different bacterial species through the lens of histogram plots. The data reveals that the most prevalent range for number of genes in bacterial species is between 3600 and 5100 genes, while the corresponding genome length is between 4 to 5.6 million base pairs. In general, the bacterial genome as suggested by the scatter plot on number of genes and genome length is not bloated.

**Figure 2:**
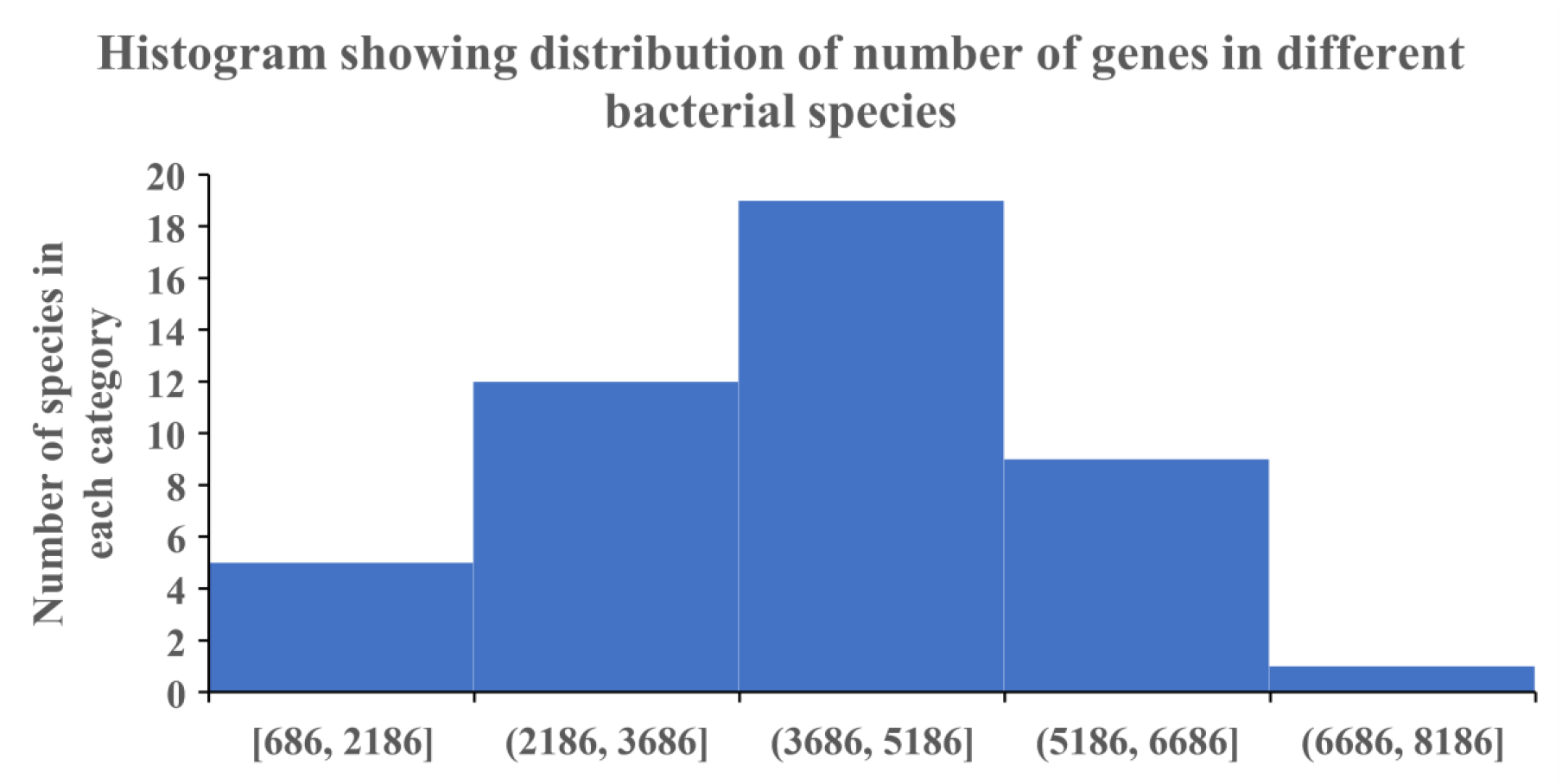
Histogram showing distribution of number of genes in different bacterial species. The data shows that most bacterial species harbour between 3600 to 5100 genes in their genome, making this gene number range most prevalent in the bacteria domain.

**Figure 3:**
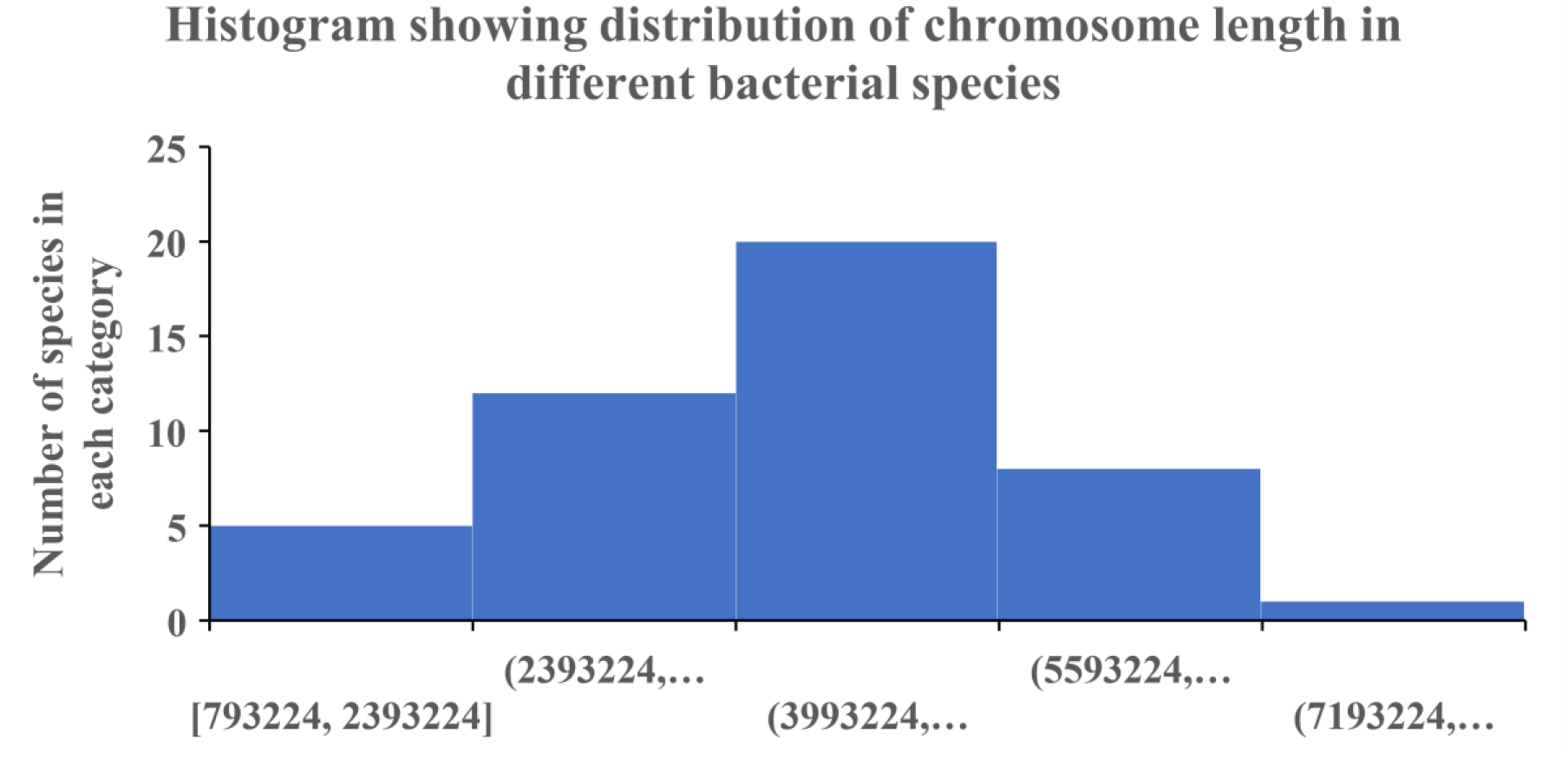
Histogram showing distribution of chromosome length in different bacterial species. The data reveals that the largest category of bacterial species harbour a genome length of between 4 million base pair to 5.6 million base pair.

Repeating the same scatter plot analysis for archaea species revealed a corresponding positive correlation between number of genes and chromosome length in archaea species. Figure 4 shows the result of this plot. Closer inspection of the plot, however, reveals that while there is a general positive correlation between number of genes and chromosome length, many data points do not fall on the trendline. This suggests deviation from the general trend that increase in number of genes lead to longer genome. Such deviations suggest repetitive sequence may expand the size of a genome. However, existence of a good correlation meant that repetitive sequence is only a nascent phenomenon in archaeal species, and may be a hallmark feature of the archaeal domain. On the evolutionary level, the trendline elucidated in this work could be further evidence that archaea is the ancestral species of modern eukaryotes.

**Figure 4:**
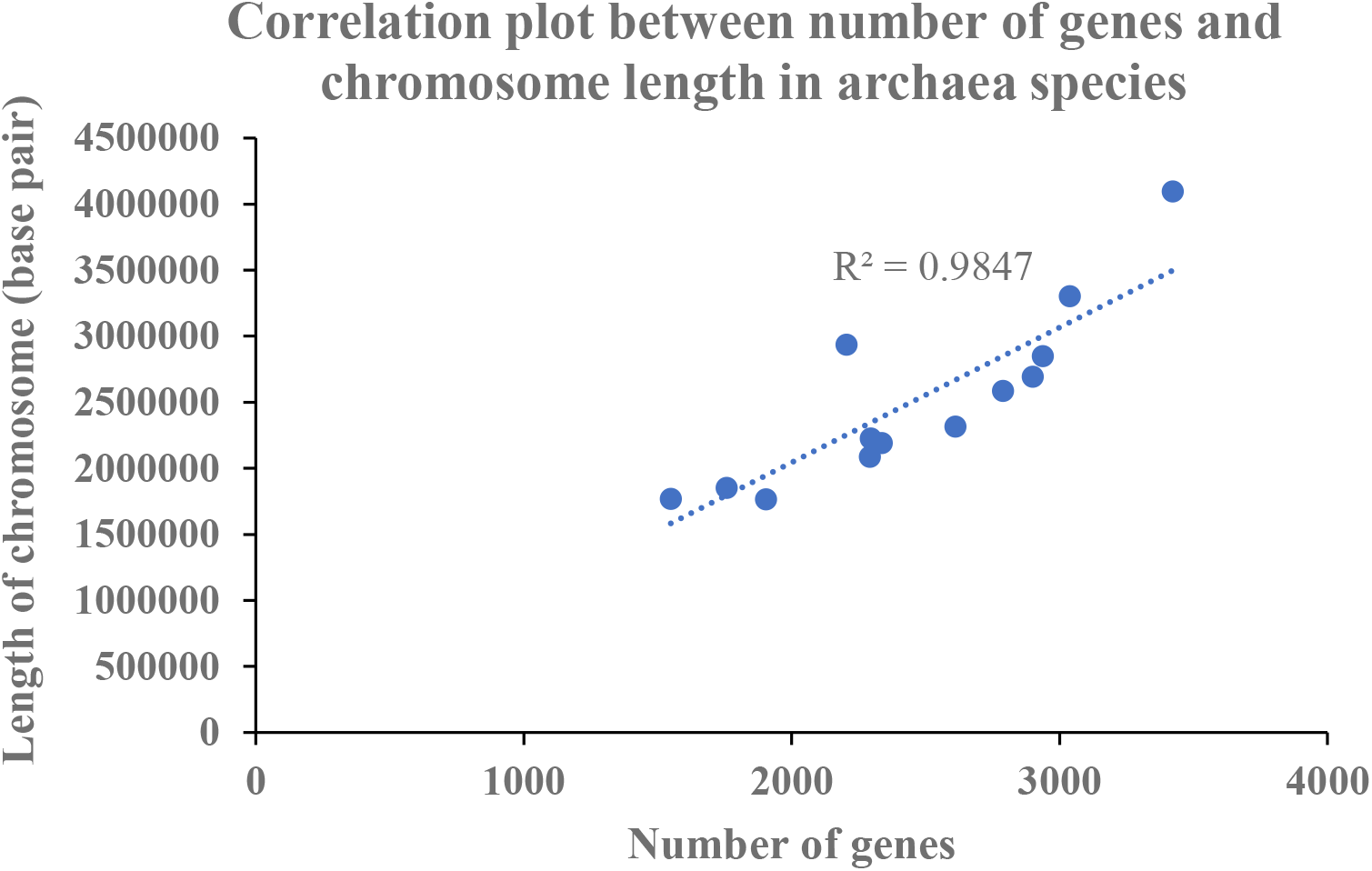
Correlation plot between number of genes and chromosome length in different archaea species. The data shows a positive correlation between number of genes and chromosome length in archaea species with a few outlier data points, implying that repetitive sequences have become more prevalent in archaea in what is a more eukaryotic-like organism.

Figure 5 and 6 shows the histogram analysis of number of genes and chromosome length distribution in archaeal species. Although the number of archaeal species profiled is much smaller than that of bacterial species, the data obtained nevertheless remain informative. Specifically, the data reveals that the most prevalent range for number of genes in archaea is 1500 to 2400 genes, which is significantly smaller than that of bacterial species. This suggests that archaeal species may be more primitive with respect to the evolution of life on Earth. On the other hand, histogram data also shows that the most prevalent range for genome length in the archaeal domain is 1.8 to 2.8 million base pair which is much shorter than that in bacterial domain. Overall, the data casts a picture where archaea and bacteria may be two independent evolutionary lineages with archaea having evolved earlier and remains more primitive and specialised compared to bacterial species. The second point is important because the more generalist bacterial species need more genes and chromosome length to genetically encode more specialised and diversified functions to colonize more habitats and niches across Earth.

**Figure 5:**
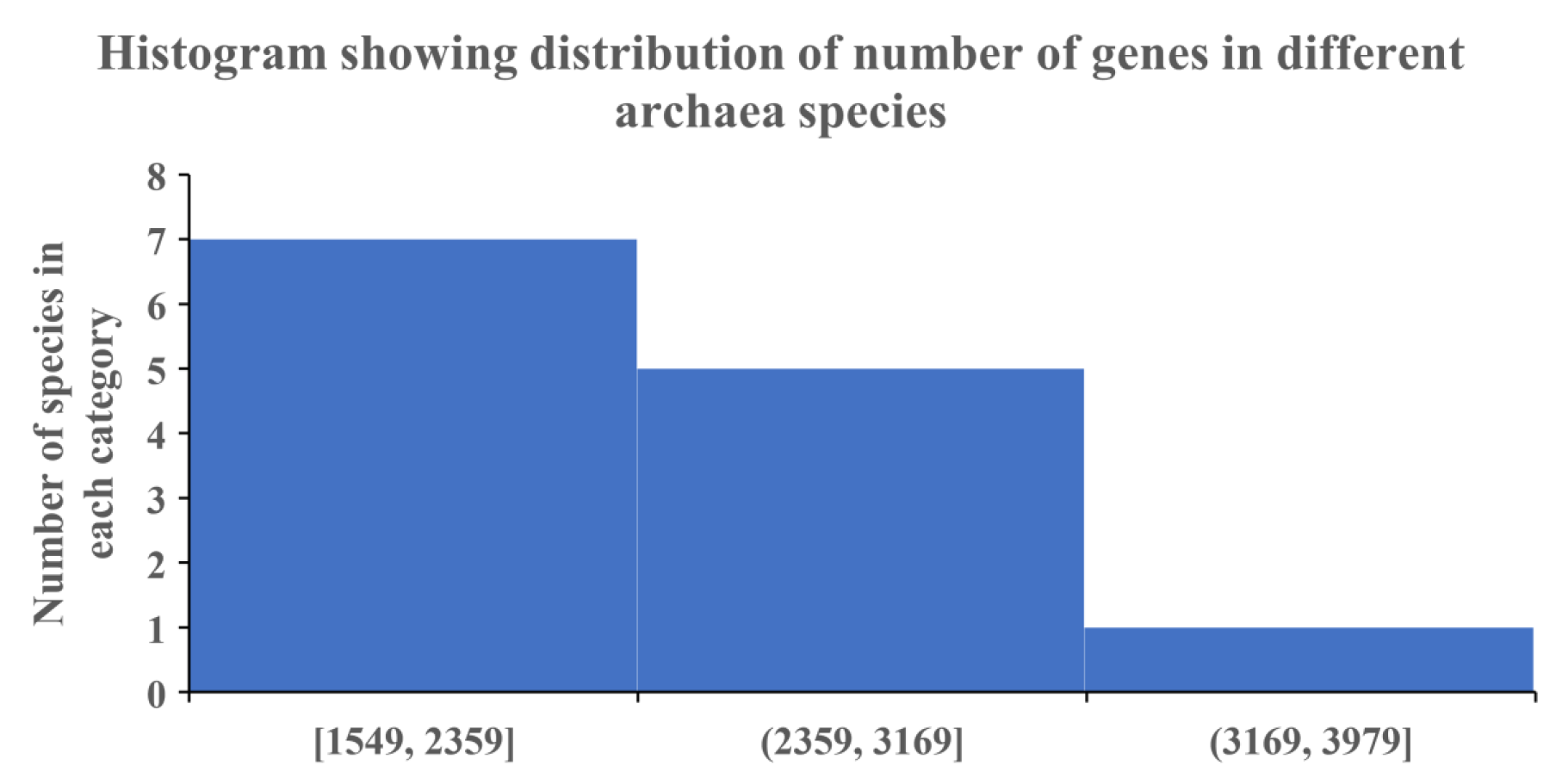
Histogram showing distribution of number of genes in different archaea species. The data shows that the most prevalent category is archaeal species with 1500 to 2400 genes in their genome.

**Figure 6:**
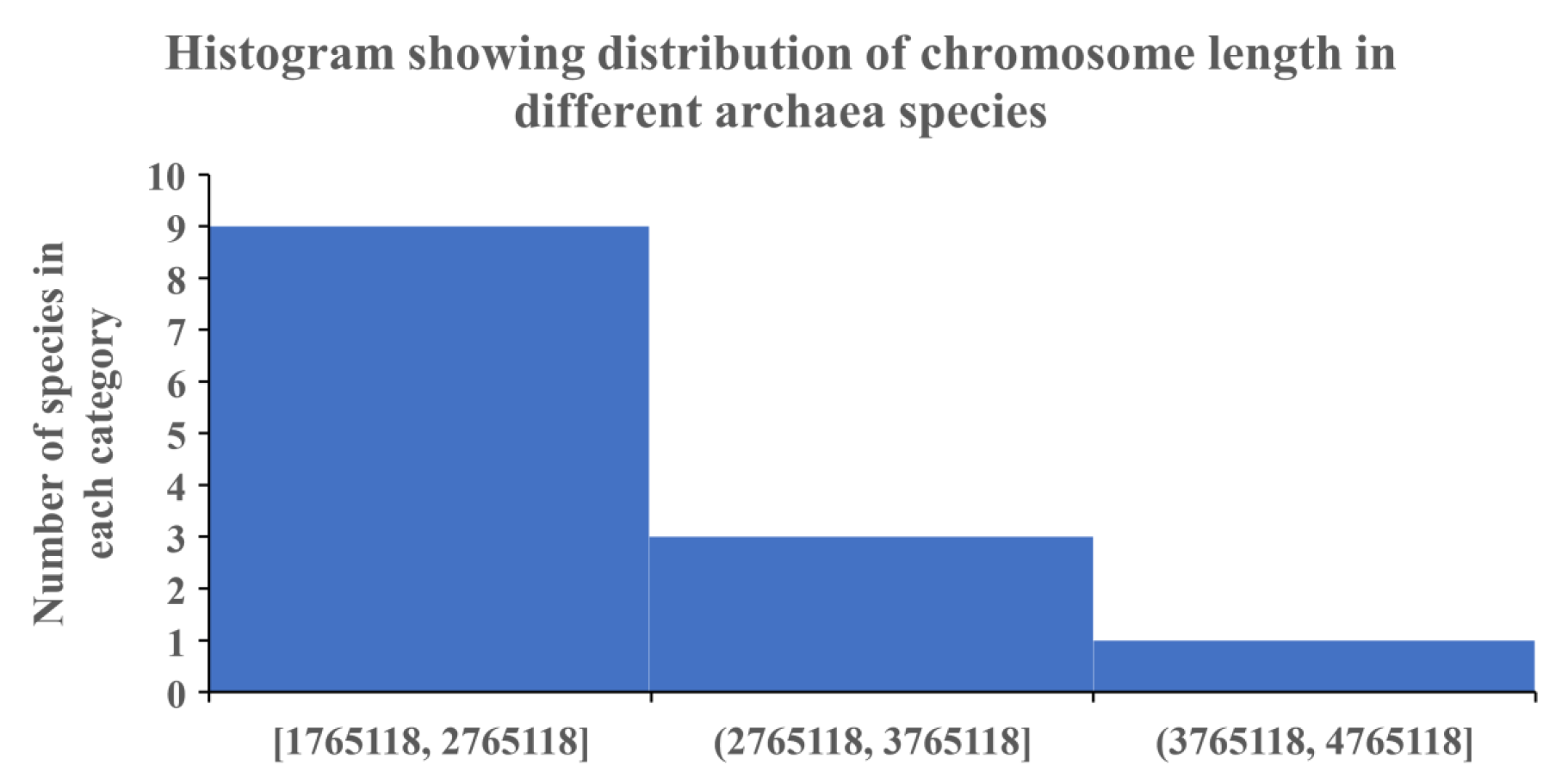
Histogram showing distribution of chromosome length in different archaea species. The data shows that a large number of archaeal species sport small genomes of length between 1.8 million base pair to 2.8 million base pair, which is significantly smaller than that of bacterial species.

Different lines of evidence suggests that archaea are the first form of cellular life on Earth [5]. Evidence includes the presence of different types of extremophiles in this domain suited for life in extreme environments prevalent on early Earth [6] [7]. In addition, the number of genes and chromosome length of archaeal species are also smaller than corresponding values in bacterial species. Given that repetitive sequences are prevalent in archaeal species and not bacterial species, it is possible that Archaea and Bacteria evolve as independent domains of life with the archaea giving rise to eukaryotes, and Bacteria continuing on in its independent evolution. Bacteria later give rise to plant cells which have less repetitive sequences. Thus, early Earth may have two different types of cellular life: Archaea and Bacteria, which have their own independent evolutionary trajectories. There may be some exchanges of genes and plasmid between the two domains along the way, but these two domains should remain fairly independent in evolution.

## Conclusions

Understanding the architecture of genomes in bacteria and archaea requires large amount of genomic data. Thanks to the proliferation of sequencing initiatives around the world, the world now has a huge and rich vein of information on the genomes of varied bacteria and archaea of different genome length. Tapping on this data resources using the system biology approach, this work reveals a strong positive correlation between number of genes and chromosome length in bacterial species, and less so in archaeal species. Overall, bacterial genome contains more genes and is longer than archaeal species. This suggests that repetitive sequence may be a nascent phenomenon in archaeal genome, and provides evidence supporting that archaeal and bacterial species follow two independent lines of evolution, with archaeal species possibly ancestral to bacterial species. In addition, the data also corroborates observations that bacterial species are more generalist in metabolism and physiology compared to archaeal species where more genes and greater chromosome length can be useful for encoding more functions.

## References

[1] K. Michalska and A. Joachimiak, “Structural genomics and the Protein Data Bank,” J. Biol. Chem., vol. 296, Jan. 2021, doi: 10.1016/j.jbc.2021.100747.

[2] S. Srinivasan et al., “Structural Genomics of SARS-CoV-2 Indicates Evolutionary Conserved Functional Regions of Viral Proteins,” Viruses, vol. 12, no. 4, p. E360, Mar. 2020, doi: 10.3390/v12040360.

[3] D. J. Downes et al., “High-resolution targeted 3C interrogation of cis-regulatory element organization at genome-wide scale,” Nat. Commun., vol. 12, no. 1, Art. no. 1, Jan. 2021, doi: 10.1038/s41467-020-20809-6.

[4] O. V. Bylino, A. N. Ibragimov, A. E. Pravednikova, and Y. V. Shidlovskii, “Investigation of the Basic Steps in the Chromosome Conformation Capture Procedure,” Front. Genet., vol. 12, 2021, Accessed: Aug. 09, 2022. [Online]. Available: https://www.frontiersin.org/articles/10.3389/fgene.2021.733937

[5] G. Caetano-Anollés et al., “Archaea: The First Domain of Diversified Life,” Archaea, vol. 2014, p. e590214, Jun. 2014, doi: 10.1155/2014/590214.

[6] N. Merino et al., “Living at the Extremes: Extremophiles and the Limits of Life in a Planetary Context,” Front. Microbiol., vol. 0, 2019, doi: 10.3389/fmicb.2019.00780.

[7] C. J. Reed, H. Lewis, E. Trejo, V. Winston, and C. Evilia, “Protein Adaptations in Archaeal Extremophiles,” Archaea, vol. 2013, p. e373275, Sep. 2013, doi: 10.1155/2013/373275.

